# Proteomic profiling reveals pleiotropic antimetabolite activity of triciribine in acute lymphoblastic leukemia

**DOI:** 10.64898/2026.03.13.710465

**Authors:** Xuekang Qi, Georgios Mermelekas, Ondrej Hodek, Luay Aswad, Quentin Glaziou, Isabelle Rose Leo, Nona Struyf, Ioannis Siavelis, Henri Colyn Bwanika, Owen F.J. Hovey, Martin Haraldsson, Annika Johansson, Tom Erkers, Katja Pokrovskaja Tamm, Janne Lehtiö, Rozbeh Jafari

## Abstract

Acute lymphoblastic leukemia (ALL) exhibits marked genetic and metabolic heterogeneity that limits the efficacy of targeted therapies. Antimetabolite strategies remain central to ALL treatment, yet the mechanisms underlying differential drug sensitivity are incompletely defined. Here, we investigated the activity of the purine analog triciribine (TCN) across diverse ALL cellular models. We find that TCN exerts potent cytotoxic effects in multiple ALL cell lines, exceeding those observed with canonical Akt inhibitors in our datasets. Phosphoproteomic analyses indicate that, at early time points, TCN does not primarily suppress Akt signaling but instead induces transient pathway activation accompanied by inhibition of cyclin-dependent kinases in both sensitive and resistant cells. The monophosphorylated metabolite TCN-P represents the predominant intracellular species and requires adenosine kinase (ADK) for activity, with ADK protein levels positively correlating with TCN sensitivity in both cell lines and primary patient samples. Using Proteome Integral Solubility Alteration profiling, we identify candidate protein interactions of TCN-P distributed across multiple cellular pathways, including nucleotide metabolism, DNA replication, and protein synthesis. These interactions are accompanied by impaired purine biosynthesis, DNA damage, translational stress, and cell-cycle arrest, as supported by time-course quantitative proteomics and immunoblot analyses. Together, these findings characterize triciribine as an antileukemic agent with pleiotropic antimetabolite activity in ALL and highlight ADK-dependent metabolism as a key determinant of therapeutic sensitivity, suggesting that ADK levels may serve as a predictive biomarker to stratify patients for triciribine-based precision treatment strategies.

## Introduction

ALL is an aggressive hematologic malignancy that affects patients of all ages and is characterized by rapid proliferation and disease progression[1, 2]. Extensive genetic and epigenetic heterogeneity underlies substantial diversity in molecular features, cellular behavior, and clinical responses to therapy. Although this complexity complicates the development of universally effective targeted treatments, leukemic cells remain uniformly dependent on high biosynthetic and metabolic flux, creating shared vulnerabilities across ALL subtypes. Altered cellular metabolism therefore represents not only a hallmark of ALL but also a functional dependency that may be less constrained by genotype [3, 4].

Nucleoside and nucleotide metabolism constitutes a fundamental requirement for leukemic cell survival and proliferation. Rapidly proliferating leukemic cells require sustained nucleotide synthesis to support DNA replication, RNA transcription, protein translation, and energy homeostasis, rendering these pathways particularly sensitive to pharmacologic perturbation [5, 6]. Disruption of these pathways has therefore been extensively exploited for therapeutic purposes [4, 7]. Nucleoside analogues (NAs) interfere with nucleotide metabolism by targeting key enzymatic steps or by mimicking endogenous nucleosides, thereby impairing nucleic acid synthesis and inducing cytotoxic stress in rapidly proliferating cells [4, 8, 9]. Such agents have demonstrated substantial clinical activity in multiple malignancies, including ALL.

Despite major advances in risk stratification and targeted therapy, the backbone of ALL treatment remains broad cytotoxic chemotherapy, including several nucleoside analogues and antimetabolites [1, 7]. While these agents have enabled high cure rates, they lack predictive biomarkers that inform patient-specific sensitivity or resistance remain limited [1, 7, 9]. As a result, response variability and treatment-related toxicity continue to pose significant challenges, particularly in relapsed or refractory disease. Identifying molecular features that govern sensitivity to cytotoxic and antimetabolite therapies therefore represents an important unmet need, both to refine existing treatment strategies and to guide the development of next-generation agents.

Triciribine (TCN) was originally developed as an adenosine analog, with its monophosphorylated form TCN-P representing the predominant intracellular and biologically active species [10]. Unlike most nucleoside analogues, which require sequential tri-phosphorylation and are incorporated into nucleic acids, TCN is unusual in that its mono-phosphorylated form is sufficient for biological activity suggesting alternative modes of action [8, 11, 12]. In clinical formulations, TCN-P functions as a water-soluble prodrug that is dephosphorylated extracellularly by phosphatases and ecto-phosphatases prior to cellular uptake [10]. Once inside the cell, TCN is re-phosphorylated by adenosine kinase (ADK), a step that is required for its cytotoxic activity[10, 13]. Consistent with this requirement, ADK expression has been associated with TCN sensitivity in multiple cellular contexts[14, 15].

Despite its classification as a nucleoside analogue, triciribine has frequently been described as a direct inhibitor of Akt signaling[16–18]. However, this model has failed to fully account for triciribine’s cellular activity which often exceeds that of canonical PI3K-Akt-mTOR pathway inhibitors and displays distinct response patterns across cancer models [19]. In addition, TCN has been associated with inhibition of DNA synthesis, RNA synthesis, protein synthesis and perturbation of purine nucleotide biosynthesis, indicating that its cellular effects may extend beyond a single signaling axis [11, 12, 20–22]. The limited predictive value of Akt pathway in clinical trials further underscores the incomplete understanding of TCN’s mechanism of action [23].

Together, these observations position triciribine as a mechanistically anomalous compound that does not conform neatly to established categories of either classical nucleoside analogues or targeted kinase inhibitors. Given this ambiguity, an unbiased and systems-level approach is required to define the molecular processes and pathways that govern triciribine response. Proteomics based on liquid chromatography–mass spectrometry (LC–MS) provides a powerful platform to examine changes in protein abundance, post-translational modifications, proteoform diversity and protein–ligand interactions at a global scale [24–27]. In this study, we applied complementary proteomic strategies, including thermal proteome profiling approaches such as Proteome Integral Solubility Alteration (PISA), quantitative time-course proteomics, and phosphoproteomics, to characterize cellular responses to TCN treatment in ALL models [26, 28–30]. Rather than seeking a single dominant target, these approaches were used to identify the pathways and cellular processes associated with TCN sensitivity and response, providing a framework for mechanistic interpretation and biomarker development for translational applications.

## Results

### Triciribine exhibits potent antileukemic activity and a distinct response profile compared with Akt inhibitors

In our previous integrative multi-omics and drug sensitivity profiling of 43 childhood ALL cell lines, we observed that triciribine displayed a response profile distinct from canonical Akt inhibitors (**Fig. 1A**) [31, 32]. Pairwise correlation analysis of drug sensitivity profiles revealed minimal correlation between triciribine and afuresertib, ipatasertib, or uprosertib (**Supplementary Fig. 1A**), whereas the shared mechanism was supported by robust mutual correlations in these inhibitors (**Supplementary Fig. 1B**). Compared with normal bone marrow samples, TCN induced cytotoxicity across the majority of ALL cell lines and subtypes, with a substantial fraction displaying hypersensitivity (selective drug sensitivity score, sDSS ≥ 8). In contrast, most PI3K–Akt–mTOR inhibitors showed markedly lower activity, with sDSS values below this threshold in the majority of ALL models (**Fig. 1B**). Consistent with these findings, pairwise correlation and dimensionality reduction analyses demonstrated generally low similarity between triciribine and other PI3K-Akt-mTOR inhibitors, with only modest correlations observed for selected PI3K or PI3K/mTOR inhibitors such as pictilisib and GDC-0084 (**Fig. 1C**).

**Figure 1.**
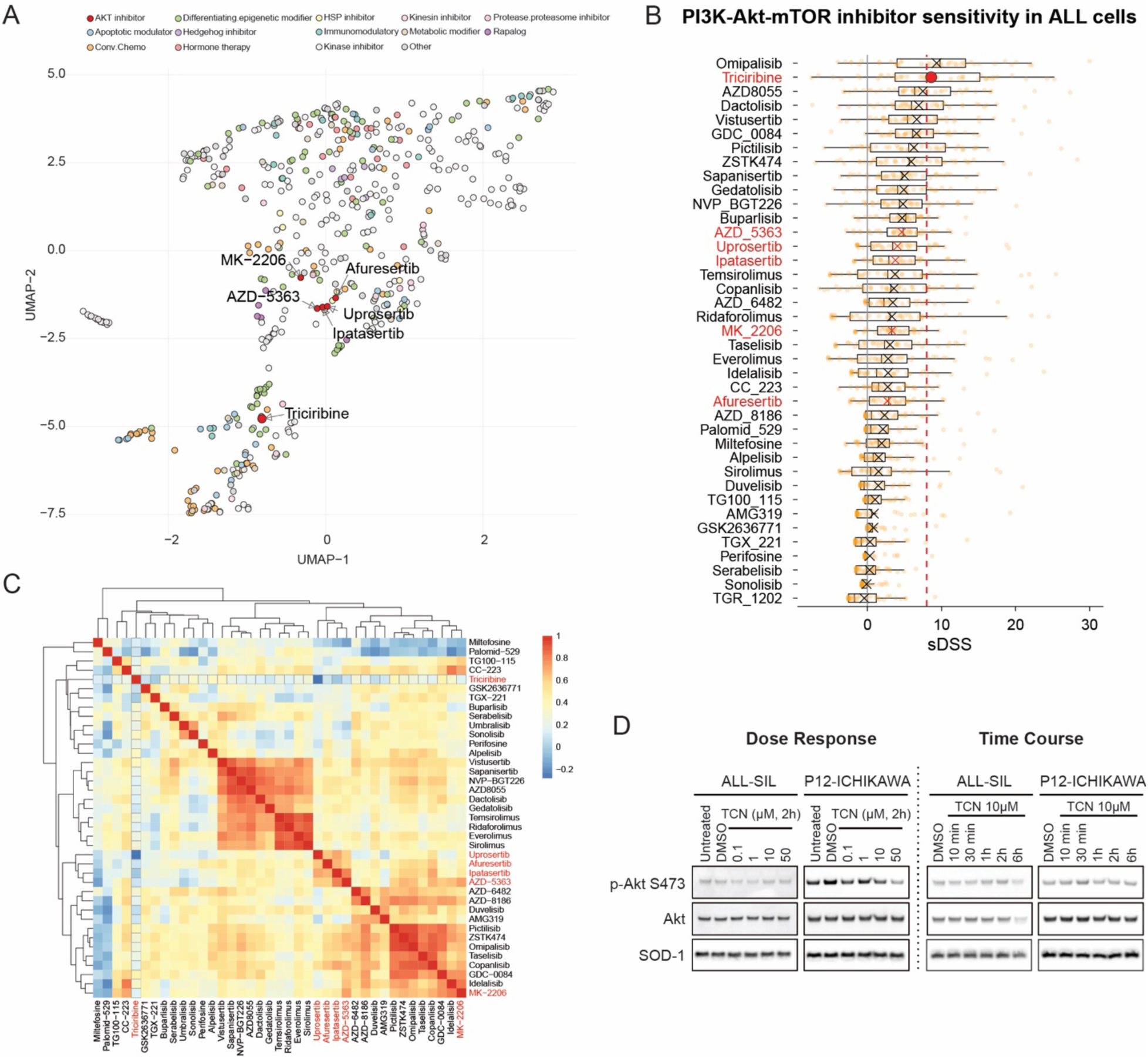
Drug sensitivity profiling of triciribine across ALL cell lines. **A**, UMAP visualization of drug sensitivity profiles for 528 compounds tested across 43 ALL cell lines. Each point represents a compound, positioned based on similarity of killing profiles. **B**, sDSS values for triciribine and PI3K-Akt-mTOR pathway inhibitors across 43 ALL cell lines. The dashed line indicates sDSS = 8. The mean sDSS value for each inhibitor is indicated by an ‘×’ or a larger dot. Akt inhibitors are indicated in red. **C**, Pairwise Spearman correlation analysis of drug sensitivity profiles for PI3K-Akt-mTOR pathway inhibitors across 43 ALL cell lines. Akt inhibitors are indicated in red. **D**, Immunoblot analysis of phosphorylated Akt (Ser473), total Akt, and the loading control SOD1 in ALL-SIL and P12-ICHIKAWA cells following treatment with indicated concentrations or durations of triciribine.

Analysis of the PRISM drug repurposing dataset [33], encompassing 4,518 compounds tested across 578 cancer cell lines representing many lineages, further validated these observations. Triciribine again segregated from the PI3K-Akt-mTOR inhibitor cluster (**Supplementary Fig. 1C**), suggesting that its distinct response profile is neither restricted to hematologic malignancies nor specific to a single dataset.

To further examine the relationship between triciribine activity and Akt signaling, we analyzed the PTEN-deficient T-ALL cell line P12-ICHIKAWA, which exhibits constitutive Akt activation [34].

Despite this phenotype, P12-ICHIKAWA cells were not hypersensitive to triciribine (sDSS = 2.8). Dose-response western blotting revealed that phosphorylation of Akt at Ser473 decreased only at higher triciribine concentrations (**Fig. 1D**). Moreover, time-course analysis demonstrated transient Akt phosphorylation following triciribine exposure in both P12-ICHIKAWA cells and the triciribine-sensitive ALL-SIL cell line (sDSS = 19.5) (**Fig. 1D**).

Together, these data indicate that triciribine exhibits potent antileukemic activity with a response profile distinct from that of canonical Akt inhibitors, suggesting that additional mechanisms may contribute to its cellular effects.

### Phosphoproteomic profiling reveals early inhibition of CDK activity and activation of mTOR and MAPK signaling following triciribine treatment

To systematically characterize kinase signaling responses to TCN, we performed time-course phosphoproteomic analyses in 4 ALL cell lines representing different lineages, immunophenotypes and sensitivities: ALL-SIL (T-ALL, ABL1 rearranged, sDSS = 19.5), P12-ICHIKAWA (T-ALL, sDSS = 2.8), KASUMI-10 (BCP-ALL, KMT2A rearranged, sDSS = 24.2), and KASUMI-2 (BCP-ALL, TCF3 rearranged, sDSS = –4.7) (**Fig. 2A**). ALL-SIL and KASUMI-

**Figure 2.**
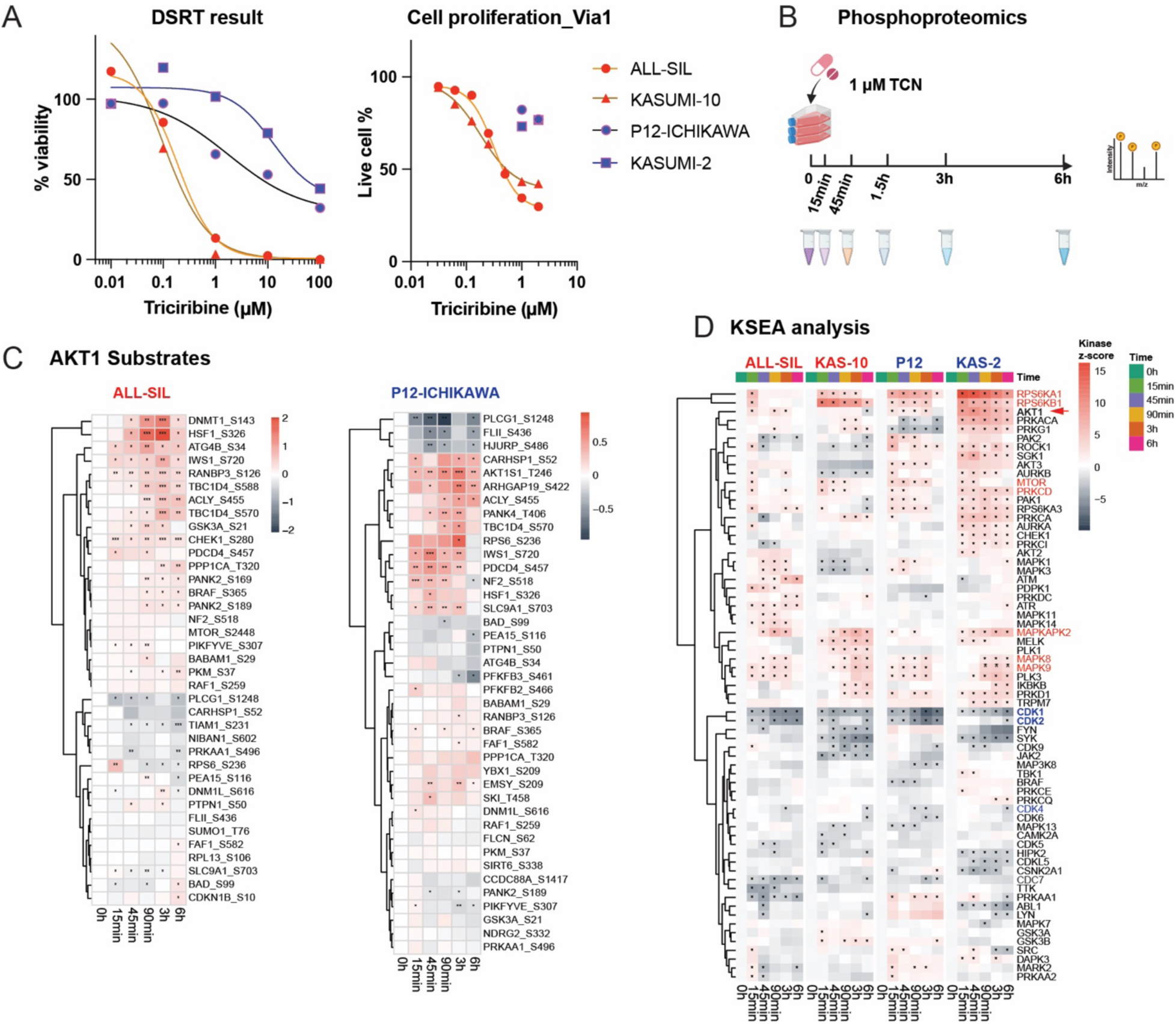
Phosphoproteomic analysis of signaling responses following triciribine treatment. **A**, Drug sensitivity testing (DSRT) and in-house cell proliferation assays comparing triciribine sensitivity in four ALL cell lines. Cells were treated with increasing concentrations of triciribine for 72 h, and cell number and viability were measured using CellTiter-Glo or Via1 cassette-NucleoCounter. **B**, Experimental design and workflow for time-course phosphoproteomic analysis. Four ALL cell lines were treated with 1 μM triciribine for the indicated time points prior to phosphopeptide enrichment and LC–MS/MS analysis. **C**, Phosphorylation changes of known Akt1 substrates across time points relative to 0 h in ALL-SIL and P12-ICHIKAWA cells. Statistical comparisons were performed using a t-test where applicable. **D**, Heatmap of kinase activity scores derived from KSEA across all time points relative to 0 h in four ALL cell lines (ALL-SIL, P12-ICHIKAWA [P12], KASUMI-10 [KAS-10], and KASUMI- 2 [KAS-2]). Kinases with p < 0.05 are marked with an asterisk. The same abbreviations are used throughout the figures.

10 were classified as sensitive, whereas P12-ICHIKAWA and KASUMI-2 showed intermediate or resistant responses. The high-throughput drug screening results were consistent with independent methods of viability measurement, despite higher apparent IC50 values and methodological differences between assays (**Fig. 2A**).

Based on IC50 values below or near 1 µM in sensitive lines, we selected 1 µM TCN for phosphoproteomic analyses. Samples were collected at baseline and at 15 min, 45 min, 1.5 h, 3 h, and 6 h after treatment, with three biological replicates per time point (**Fig. 2B**). Principal component analysis demonstrated clear temporal separation following triciribine exposure, with tight clustering of replicates, indicating robust phosphoproteomic measurements (**Supplementary Fig. 2A**). In contrast, global protein abundance showed minimal changes during the early time points (**Supplementary Fig. 2B**), supporting independent analysis of phosphoproteomic data.

Analysis of known Akt substrates, curated from PhosphoSitePlus, revealed that phosphorylation of the majority of Akt substrates increased following triciribine treatment across all four cell lines, despite variability in individual substrate dynamics (**Fig. 2C and Supplementary Fig. 2C**). Phosphorylation levels typically peaked between 30 min and 3 h, consistent with immunoblotting results (**Fig. 1D**).

Kinase–substrate enrichment analysis (KSEA) was performed using baseline samples as controls [35]. Across all 4 cell lines, cell-cycle–associated kinases CDK1, CDK2, and CDK4, were consistently inhibited following TCN treatment (**Fig. 2D**). In contrast, kinases within the mTOR pathway (mTOR, RPS6KB1) and MAPK signaling pathways (MAPKAPK2, MAPK8/9, RPS6KA1) were consistently activated. Akt1 activity was transiently increased in all 4 cell lines at early time points, with reduced activity observed at later time points in selected models.

Overall, these results demonstrate that triciribine induces reproducible phosphoproteomic responses across ALL cell lines, characterized by early CDK inhibition and activation of mTOR and MAPK signaling, independent of cellular sensitivity.

### PISA profiling identifies candidate triciribine-interacting proteins across the proteome

Given the limited early inhibition of Akt signaling, we next sought to identify candidate direct protein interactions of triciribine using PISA profiling [29]. Prior studies have shown that triciribine is rapidly transported into cells and phosphorylated by ADK to generate the predominant intracellular species TCN-P, resulting in substantial intracellular accumulation (**Fig. 3A**) [12].

**Figure 3.**
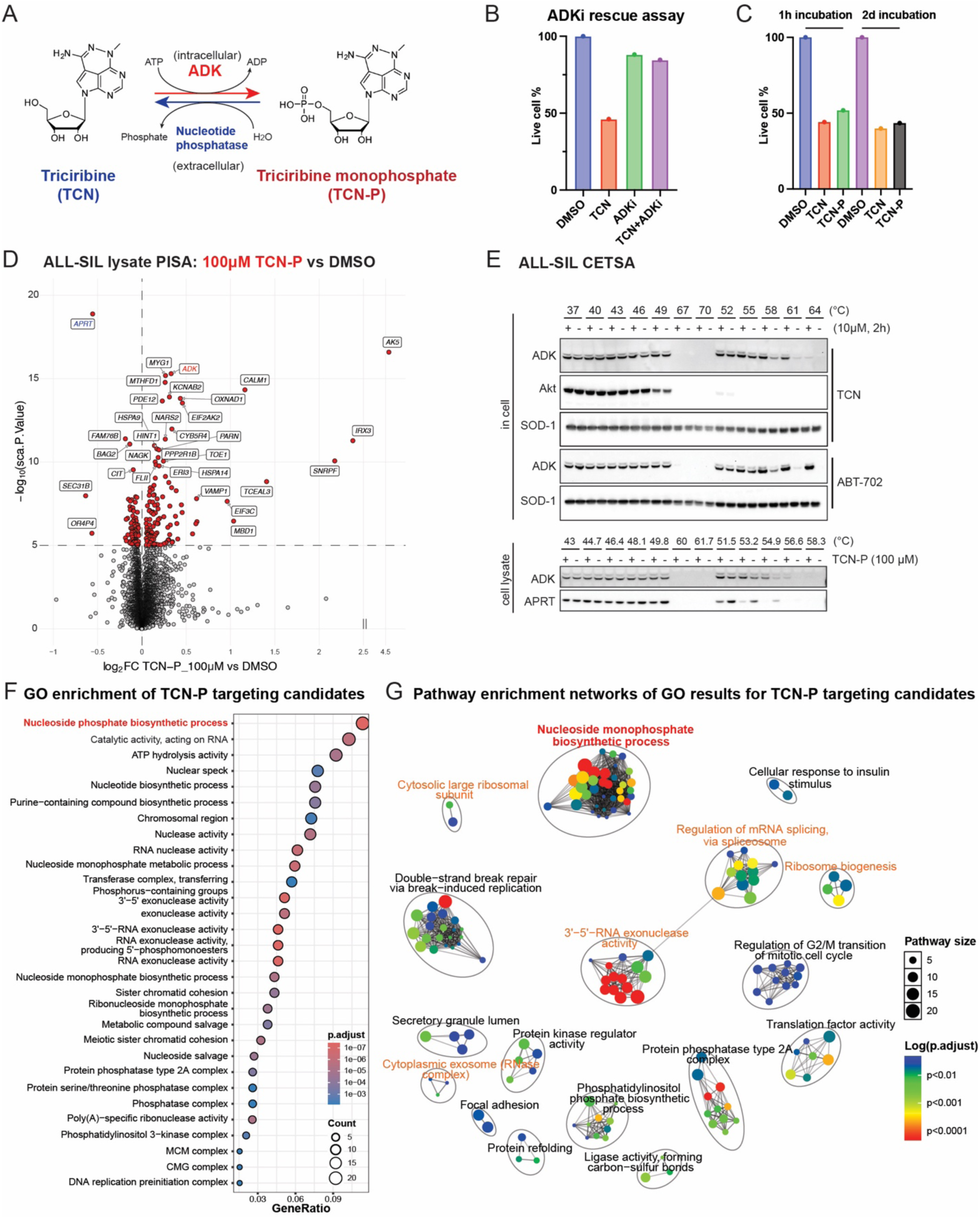
Identification of candidate triciribine-interacting proteins by PISA profiling. **A**, Chemical structures of TCN and TCN-P, and schematic representation of their interconversion by ADK and nucleotide phosphatases. ATP, Adenosine Triphosphate; ADP, Adenosine Diphosphate. **B**, Cell viability and cell number measurements in ALL-SIL cells treated with triciribine (4 μM), the ADK inhibitor ABT-702 (5 μM), or their combination for 1 h followed by culture in drug-free medium for 48 h. **C**, Cell viability and cell number measurements in ALL-SIL cells following short-term (1 h) or continuous (48 h) exposure to triciribine or TCN-P. **D**, Volcano plot of PISA results showing changes in protein solubility following treatment with 100 μM TCN-P compared with DMSO control. **E**, Cellular and lysate-based CETSA immunoblots demonstrating thermal stabilization of ADK following treatment with triciribine or TCN-P. **F**, Gene Ontology (GO) enrichment analysis of candidate TCN-P–interacting proteins identified by PISA. **G**, Pathway enrichment network visualization of GO terms enriched among candidate TCN-P–interacting proteins.

We confirmed that TCN-P is the dominant intracellular form of triciribine in ALL cells. Pharmacologic inhibition of ADK using ABT-702 fully rescued triciribine-induced cytotoxicity in ALL-SIL cells (**Fig. 3B**), confirming the requirement of ADK-mediated phosphorylation. Short-term exposure to triciribine or TCN-P was sufficient to induce cytotoxicity comparable to continuous treatment, indicating rapid uptake and conversion to TCN-P (**Fig. 3C**).

Based on published intracellular concentration estimates [12], PISA profiling was performed using ALL-SIL lysate treated with 100 µM TCN-P to approximate intracellular exposure corresponding to 1 µM extracellular triciribine. For comparison, 100 µM TCN and 1 µM TCN-P were included as control conditions. ADK served as a positive control, and four biological replicates were analyzed per condition (**Supplementary Fig. 3A**).

Relative to vehicle control, 100 µM TCN-P yielded the largest number of candidate interacting proteins (**Fig. 3D and Supplementary Fig. 3B,C**). ADK emerged as the top hit for both TCN and TCN-P, consistent with in-cell and lysate CETSA (cellular thermal shift assay) validation (**Fig. 3E and Supplementary Fig. 3D**). In contrast, no significant thermal stabilization of Akt was observed (**Fig. 3E and Supplementary Fig. 3E**), whereas multiple components of the Akt regulator PI3K and PP2A complexes showed altered stability following TCN-P treatment (**Supplementary Fig. 3F,G**).

To define high-confidence candidate interactions, we applied a stringent DEqMS scaled p-value threshold (sca.P.value < 10⁻⁵), yielding 199 candidate TCN-P-interacting proteins with at least one identified peptide (**Fig. 3D**). Gene ontology analysis revealed enrichment across diverse biological processes, with nucleoside phosphate biosynthetic processes representing the most significant cluster, along with DNA replication, RNA-associated processes, protein translation, protein folding, cell cycle related processes (**Fig. 3F,G**).

### TCN-P engages multiple metabolic and biosynthetic pathways and induces broad cellular stress responses in ALL-SIL cells

Previously reported reductions in intracellular ATP and GTP pools and inhibition of nucleic acid synthesis following triciribine exposure [12] are in line with our PISA findings, which identified stability shifts in multiple proteins involved in de novo and salvage purine biosynthesis or its regulation, including ADK, APRT, AK5, PRPS, NUDT5, ATIC, and MTHFD1 (**Fig. 4A**, **Fig. 3D and Supplementary Fig. 3D**).

**Figure 4.**
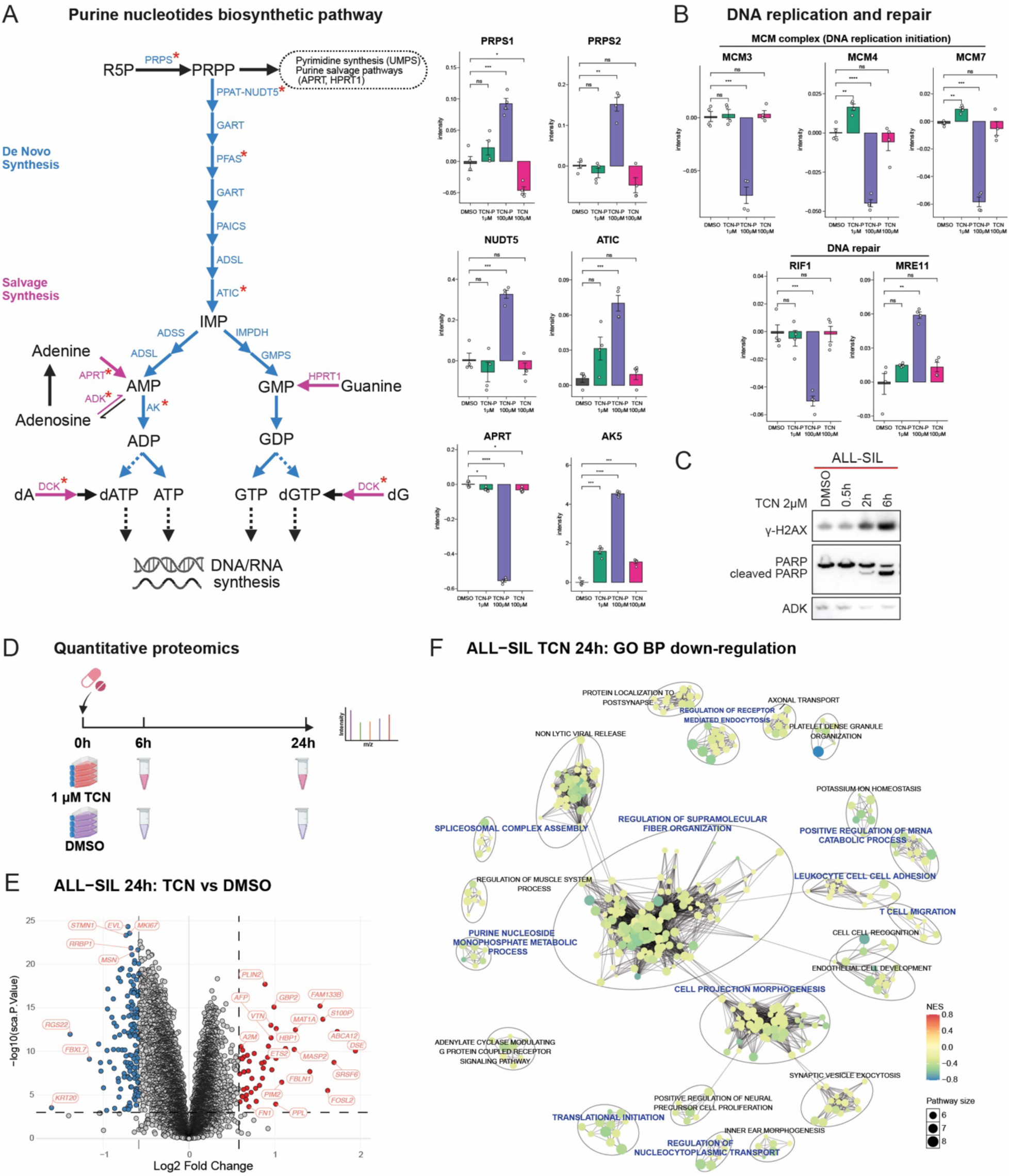
Proteomic and biochemical characterization of cellular responses to triciribine in ALL-SIL cells. **A**, Schematic overview of de novo and salvage purine nucleotide biosynthesis pathways. Enzymes identified as candidate TCN-P–interacting proteins are indicated. **B**, Bar plots showing candidate TCN-P–interacting proteins involved in DNA replication and repair. **C**, Immunoblot analysis of γ-H2AX and PARP in ALL-SIL cells following triciribine treatment. **D**, Experimental design and workflow for quantitative proteomic analysis. Four ALL cell lines were treated with DMSO or 1 μM triciribine for 6 h or 24 h prior to LC–MS/MS analysis. **E**, Volcano plot of differential protein abundance in ALL-SIL cells treated with triciribine for 24 h compared with DMSO. **F**, Visualization of significantly downregulated biological process networks identified by ssGSEA in triciribine-treated ALL-SIL cells.

In addition to purine metabolism, candidate TCN-P interactions were identified among proteins involved in DNA replication and repair [36, 37] (**Fig. 4B**). While TCN-P is not incorporated into nucleic acids, these interactions suggest that its cellular effects may extend beyond direct metabolic inhibition to broader biosynthetic and stress-related processes. Consistent with this notion, triciribine treatment induced DNA damage at early time points, as evidenced by γ-H2AX induction and PARP cleavage, preceding detectable changes in cell proliferation or viability (**Fig. 4C**).

To assess the downstream consequences of this broad interaction landscape, we performed time-course quantitative proteomics in ALL-SIL cells at 6 h and 24 h following triciribine treatment (**Fig. 4D**). Although relatively few proteins exhibited large changes in abundance at these early time points (**Fig. 4E**), pathway-level analyses revealed coordinated alterations across multiple biological processes. Single-sample GSEA was consistent with suppression of purine biosynthesis and protein translation, alongside induction of DNA damage response pathways, proteotoxic stress responses, and alterations in cytoskeletal organization and intracellular transport (**Fig. 4F and Supplementary Fig. 4A,B**).

Collectively, these findings indicate that TCN-P engages a diverse set of cellular proteins and pathways, resulting in broad metabolic and biosynthetic stress rather than disruption of a single dominant target or process. In ALL-SIL cells, triciribine exposure is therefore associated with a pleiotropic cellular response consistent with antimetabolite activity.

### ADK abundance correlates with triciribine sensitivity, and comparative proteomics across ALL models highlights sensitivity-associated response programs

Given the broad interaction profile of TCN-P, we next examined whether the abundance of candidate interacting proteins correlated with triciribine sensitivity across ALL models. Pairwise correlation analysis of baseline protein levels and triciribine drug sensitivity across 43 ALL cell lines identified ADK as the interacting protein most strongly associated with triciribine response (**Fig. 5A,B**). This association was consistent with prior transcript-level observations and was recapitulated in the 4 cell lines selected for proteomic analyses (**Supplementary Fig. 5A and Fig. 5B**).

**Figure 5.**
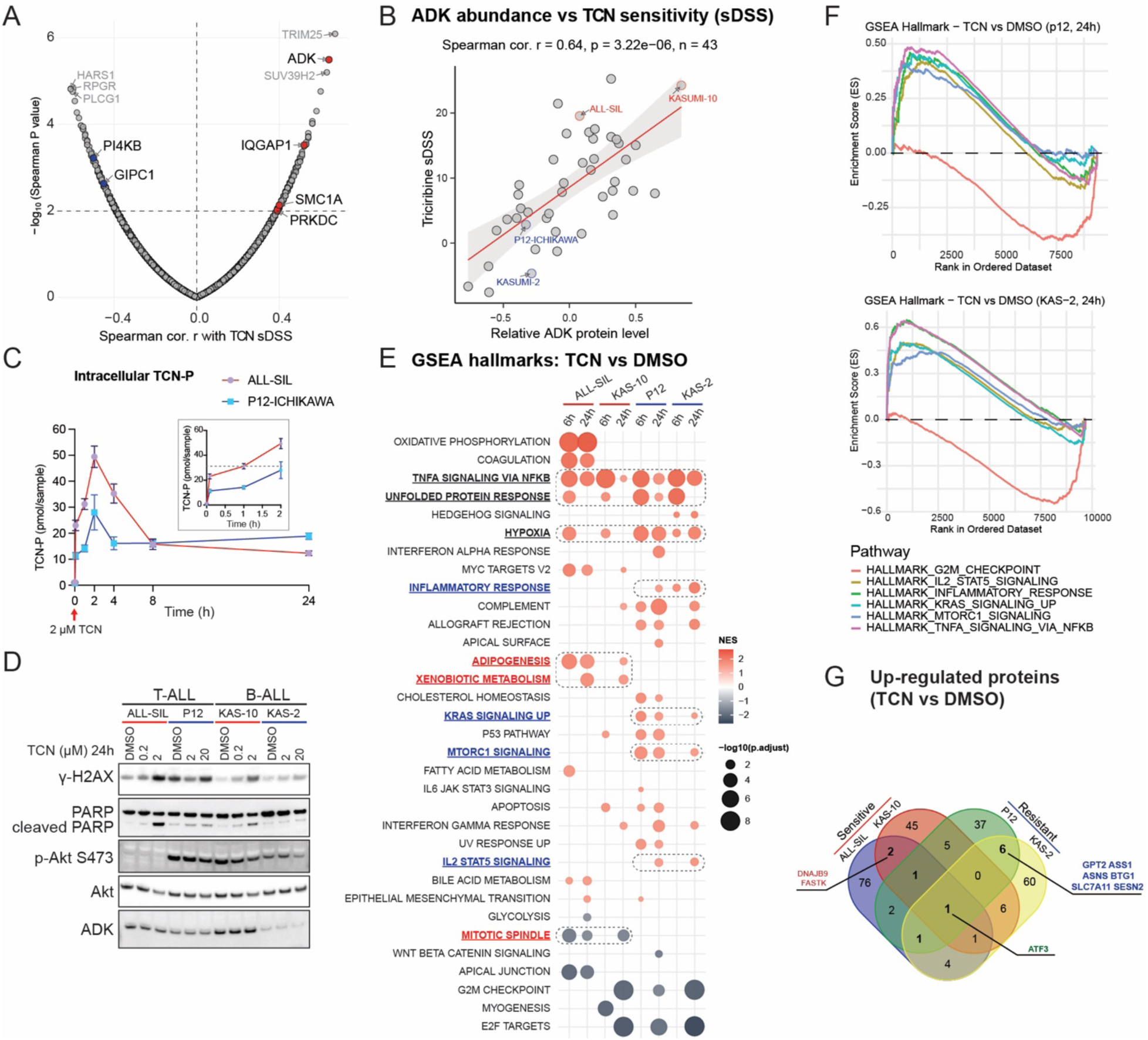
Association of ADK abundance with triciribine sensitivity and comparative proteomic responses across ALL models. **A**, Spearman correlation analysis between baseline protein abundance and triciribine sensitivity (sDSS) across ALL cell lines. Significant candidate TCN-P–interacting proteins are highlighted with black labels. **B**, Correlation between triciribine sDSS and ADK protein abundance in ALL cell lines. **C**, Time-course quantification of intracellular TCN-P concentrations in ALL-SIL and P12-ICHIKAWA cells following treatment with 2 μM triciribine. **D**, Immunoblot analysis of γ-H2AX, PARP, phosphorylated Akt (Ser473), and total Akt in ALL cell lines following treatment with increasing concentrations of triciribine. **E**, Hallmark gene set enrichment analysis (GSEA) comparing total proteome changes between triciribine- and DMSO-treated samples at 6 h and 24 h across four ALL cell lines. **F**, Representative GSEA enrichment plots of pathways consistently upregulated in less sensitive cell lines following triciribine treatment. **G**, Venn diagram showing the overlap of proteins significantly upregulated following triciribine treatment at both 6 h and 24 h across four ALL cell lines.

To determine whether ADK abundance influences intracellular TCN-P dynamics, we performed time-course LC–MS/MS quantification of triciribine and TCN-P in ALL-SIL (ADK-high, sensitive) and P12-ICHIKAWA (ADK-low, less sensitive) cells following exposure to triciribine. In both cell lines, intracellular TCN-P levels increased rapidly, peaking within approximately 2 h, and subsequently declined to a stable plateau, consistent with (partial) export of TCN-P into the extracellular compartment [38] (**Fig. 5C and Supplementary Fig. 5B**). Notably, the maximal TCN-P concentration achieved differed substantially between the two models and closely tracked ADK protein abundance, whereas steady-state TCN-P levels were comparable. The intracellular TCN-P concentration reached in ALL-SIL cells within 1 h exceeded the maximal level observed in P12-ICHIKAWA cells, supporting the notion that short-term exposure is sufficient to induce cytotoxicity in ADK-high models (**Fig. 5C and Fig. 3C**). In contrast, intracellular triciribine levels remained similar between cell lines throughout the time course (**Supplementary Fig. 5B**).

Increasing extracellular TCN concentrations in ADK-low cells failed to further enhance DNA damage or apoptotic responses (**Fig. 5D**), indicating that ADK abundance limits maximal intracellular formation of TCN-P and, consequently, the magnitude of downstream cellular stress. Because triciribine sensitivity is unlikely to be determined solely by drug activation, we further compared early proteomic response programs across ALL models with differing sensitivities. Using the same quantitative proteomics workflow applied to ALL-SIL, we analyzed TCN-induced proteomic changes in P12-ICHIKAWA, KASUMI-10, and KASUMI-2 cells. Hallmark pathway analysis revealed both shared and divergent response patterns (**Fig. 5E**). Across all models, triciribine treatment was associated with enrichment of unfolded protein response, hypoxia and NF-κB signaling pathways, consistent with generalized cellular stress. In contrast, suppression of mitotic and proliferative programs was more pronounced in sensitive models, whereas less sensitive models exhibited stronger induction of adaptive stress and survival-associated pathways, including mTORC1, KRAS-related, and inflammatory signaling (**Fig. 5F**). Several stress-associated proteins, including GPT2, SLC7A11, ASNS, and SESN2[39, 40], showed greater induction in less sensitive cells following triciribine treatment (**Fig. 5G and Supplementary Fig. 5C–E**).

Together, these results suggest that ADK abundance governs the extent of intracellular TCN-P accumulation and thereby influences the magnitude of TCN-induced cellular stress. Downstream proteomic responses vary across ALL models and reflect differential engagement of adaptive pathways rather than activation of a single resistance mechanism.

### Associations between ADK expression, pathway dependencies, and triciribine response in clinical samples

To explore clinical relevance, we assessed TCN sensitivity in primary ALL samples *ex vivo*. TCN at 1 µM achieved approximately 50% growth inhibition in 5 out of 10 samples (**Fig. 6A**). ADK mRNA levels showed a positive and statistically significant association with triciribine sensitivity (**Fig. 6B**). Consistent with the association observed in ALL samples, *ex vivo* testing in AML cohorts also showed increased triciribine sensitivity in samples with higher ADK levels. [41, 42] (**Fig. 6C**). In an AML cohort of 488 patient samples, high ADK abundance was associated with inferior prognosis in chemotherapy-treated patients, but not in untreated patients (**Supplementary Fig. 6A, B**), suggesting that TCN may represent a useful therapeutic complement to current chemotherapy regimens in AML.

**Figure 6.**
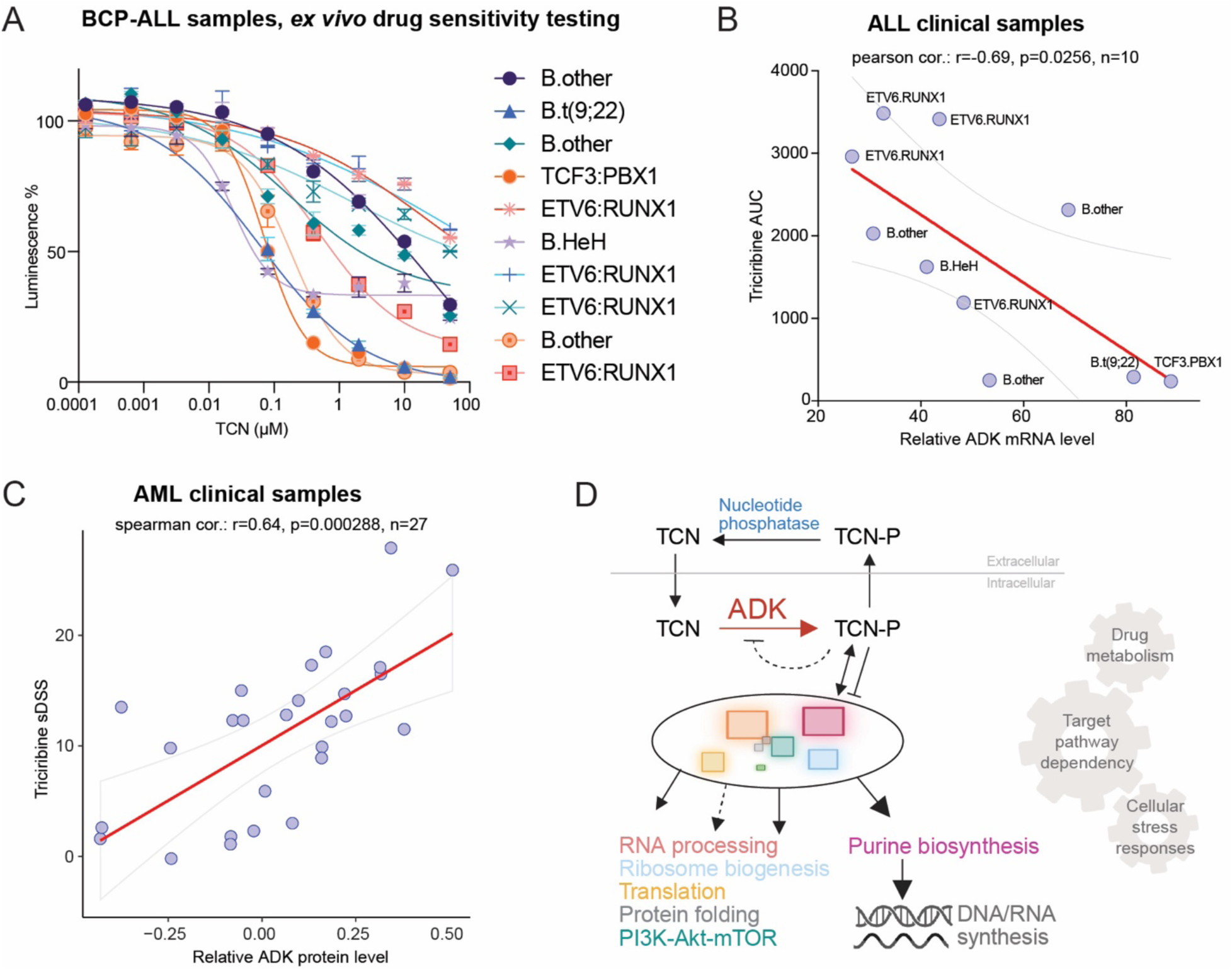
*Ex vivo* triciribine sensitivity and association with ADK expression in clinical samples. **A**, *Ex vivo* drug sensitivity testing of triciribine in primary ALL samples (n = 10) following 72 h incubation. **B**, Correlation between triciribine sensitivity (AUC, area under the curve) and ADK protein abundance in ALL samples. **C**, Correlation between triciribine sensitivity (sDSS) and ADK protein abundance in AML samples. **D**, Schematic working model summarizing proposed cellular processes associated with triciribine activity in ALL cells.

Together, these findings suggest that ADK expression, targeting pathway dependency and downstream metabolic features jointly influence cellular responses to triciribine (**Fig. 6D**).

## Discussion

In this study, we applied integrated proteomic approaches to characterize the cellular responses and candidate targets associated with triciribine treatment in acute lymphoblastic leukemia. Across multiple ALL models, triciribine exhibited potent antileukemic activity and a response profile distinct from canonical Akt inhibitors. Rather than inducing immediate suppression of Akt signaling, phosphoproteomic analyses revealed early inhibition of cyclin-dependent kinases accompanied by activation of stress-associated signaling pathways, including mTOR and MAPK, with evidence of delayed Akt modulation at later time points. Together, these findings suggest that triciribine is unlikely to function predominantly through immediate and direct suppression of Akt signaling in ALL, but instead elicits a broader cellular stress response consistent with indirect and multi-layered mechanisms of action. Later effects on Akt pathway activity may emerge downstream of cellular stress.

Using PISA profiling, we identified a diverse landscape of candidate protein interactions for the active metabolite TCN-P. These interactions were distributed across multiple metabolic and biosynthetic pathways, including nucleotide metabolism, DNA replication, protein synthesis, and RNA-associated processes. Rather than pointing to a single dominant downstream target, this interaction profile supports a model in which TCN-P perturbs multiple cellular systems simultaneously. Consistent with this interpretation, quantitative proteomics revealed coordinated pathway-level changes following triciribine exposure, including suppression of purine biosynthesis and translational programs alongside activation of DNA damage responses and proteotoxic stress pathways. These findings collectively characterize triciribine as a pleiotropic antimetabolite whose cytotoxic effects arise from broad biosynthetic disruption rather than selective inhibition of an individual enzyme or signaling node.

A key finding of this study is the identification of ADK as a key determinant of triciribine sensitivity across ALL models. ADK catalyzes the conversion of triciribine to TCN-P, the predominant intracellular and biologically active species, and ADK abundance showed the strongest correlation with drug sensitivity among candidate interacting proteins. Kinetic analyses further demonstrated that ADK levels may govern the maximal intracellular accumulation of TCN-P, thereby shaping the magnitude of downstream cellular stress. Importantly, increasing extracellular triciribine concentrations failed to overcome low ADK expression, indicating that intracellular activation rather than drug availability limits cytotoxic efficacy. These observations position ADK as a critical gatekeeper of triciribine activity and a rational biomarker for patient stratification.

Although ADK abundance strongly influences triciribine sensitivity, our data also suggest that downstream cellular context contributes to differential responses. Comparative proteomic analyses across ALL models revealed shared stress-associated responses following triciribine exposure, alongside variability in adaptive signaling programs. Less sensitive models preferentially activated pathways associated with metabolic adaptation and survival, including mTORC1, KRAS-related signaling, and inflammatory responses, whereas suppression of proliferative programs was more pronounced in sensitive cells. These findings support a model in which triciribine sensitivity reflects both efficient intracellular activation and the capacity, or failure, of leukemic cells to engage compensatory stress responses, rather than dependence on a single resistance mechanism. Our findings also have implications for the clinical development of triciribine. Although the present study primarily focuses on ALL, the association between ADK levels and TCN sensitivity appears consistent across hematologic malignancies. Previous clinical studies have reported limited objective responses despite evidence of biological activity and acceptable toxicity profiles.

Notably, intracellular accumulation of TCN-P in patient leukemic blasts has been substantially lower than that observed in cell line models [23], raising the possibility that insufficient intracellular activation, rather than intrinsic insensitivity of downstream targets, may have constrained therapeutic efficacy. Consistent with this notion, our PISA analyses showed dose-dependent TCN-P-associated changes in protein solubility profiles, suggesting that suboptimal intracellular concentrations may fail to induce the broad biosynthetic stress required for effective cytotoxicity. These observations underscore the importance of considering intracellular drug metabolism and activation, in addition to target engagement, when evaluating antimetabolite therapies.

Several limitations of this study should be acknowledged. First, while proteomic approaches provide an unbiased and systems-level view of drug responses, they cannot distinguish primary from secondary effects with certainty. Some observed proteomic changes may therefore reflect downstream consequences of metabolic disruption rather than direct TCN-P interactions. Second, the cell lysate–based PISA approach may underrepresent membrane-associated proteins and transporters, potentially missing factors involved in TCN uptake, export, or dephosphorylation that could influence intracellular drug exposure. For example, the proposed efflux transporter ABCC1/MRP1 was not identified as significant in our PISA, although it may contribute to triciribine export [43]. Third, although quantitative proteomics provided functional support for candidate interactions, comprehensive validation of individual targets through genetic or biochemical perturbation was beyond the scope of this study. Finally, *in vivo* pharmacokinetic and pharmacodynamic relationships remain to be fully defined and will be essential for translating these findings into clinical strategies.

In summary, our study provides a comprehensive proteomic characterization of TCN activity in ALL and supports a model in which TCN functions as a pleiotropic antimetabolite whose efficacy is governed primarily by intracellular activation through ADK. By defining the interaction landscape and downstream cellular responses associated with TCN-P, this work establishes a framework for biomarker-guided evaluation of triciribine and highlights the importance of drug metabolism and biosynthetic stress in shaping antileukemic responses.

## Methods

### Cell lines and cell culture

ALL-SIL, P12-ICHIKAWA, and KASUMI-2 cells were maintained in RPMI 1640 medium containing 2 mM stable glutamine (L-Ala–L-Gln dipeptide; Sigma-Aldrich), supplemented with 10% fetal bovine serum (FBS; Sigma-Aldrich), 20 mM HEPES (Gibco), 1 mM sodium pyruvate (Sigma-Aldrich), 1× MEM non-essential amino acids (Sigma-Aldrich), and 1× penicillin–streptomycin (Sigma-Aldrich), and cultured at 37°C in a humidified atmosphere with 5% CO₂. KASUMI-10 cells were maintained in Iscove’s Modified Dulbecco’s Medium (IMDM; Sigma-Aldrich) supplemented with 20% FBS and 1× penicillin–streptomycin. Cell line origin and authentication information have been reported previously [31]. All cell lines were routinely tested for mycoplasma contamination using the MycoAlert Mycoplasma Detection Kit (Lonza) and authenticated by short tandem repeat (STR) profiling (Eurofins Genomics). Total and viable cell numbers were routinely assessed using the Via1 cassette, containing the fluorophores acridine orange (AO) and 4′,6-diamidino-2-phenylindole (DAPI), in combination with the NucleoCounter system (ChemoMetec) for cell viability monitoring and proliferation assays.

### Clinical samples and drug sensitivity assay

Primary ALL and AML samples were obtained with informed consent under protocols approved by the Regional Ethical Review Board in Stockholm (2015/1998-3; 2003/02-445; 2018/38-32). All experiments involving patient material were performed *ex vivo*, and no clinical interventions were conducted.

For triciribine sensitivity testing, cells were resuspended in Hematopoietic Progenitor Expansion Medium DXF supplemented with Cytokine Mix E (Merck; C-28021, C-39891) and seeded at 1 × 10⁶ cells/mL in 96-well flat-bottom white plates (Thermo Fisher Scientific; Cat. No. 136101). DMSO levels were normalized across the plate with a maximum final concentration of 0.1%. Triciribine was serially diluted to final concentrations ranging from 0.0001 to 50 µM in a 96-well plate format.

Cells were incubated with triciribine for 72 h, after which cell viability was measured using CellTiter-Glo 2.0 ATP reagent (Promega; G9242) on a Victor 3V plate reader. Relative cell viability values were subsequently curve-fitted using a nonlinear regression model (four-parameter sigmoidal model) in GraphPad Prism to determine area under the curve (AUC) values where applicable.

### Reagents and antibodies

Triciribine (A8541) and ABT-702 (C5282) were purchased from ApexBio Technology. Triciribine monophosphate (TCN-P; 37985) was obtained from Cayman Chemical. Primary antibodies included ADK (AM8619b-ev; Nordic Biosite), SOD1 (sc-17767; Santa Cruz Biotechnology), PARP (9532), Akt (4691), and phospho-Akt (Ser473; 4060) (Cell Signaling Technology). LI-COR IRDye secondary antibodies were used for detection.

### In-cell CETSA

In-cell CETSA was performed as described previously [31, 44]. ALL-SIL cells (1.0 × 10⁶ cells/mL) were treated with triciribine (10 μM), ABT-702 (10 μM), or DMSO for 2 h at 37°C. Cells were aliquoted and subjected to heating across a temperature gradient (37-70°C) for 3 min, followed by cooling and snap-freezing. Cell lysates were prepared by repeated freeze–thaw cycles, clarified by centrifugation, and analyzed by immunoblotting.

### Cell lysate CETSA

ALL-SIL cell lysates were prepared by freeze–thaw lysis in 100 mM HEPES containing 20 mM MgCl₂. Cleared lysates were incubated with 100 μM TCN-P or DMSO for 5 min at room temperature prior to thermal treatment as described above.

### Sample preparation for mass spectrometry

PISA sample preparation was performed as described previously [29]. ALL-SIL cell lysates were treated with DMSO, 1 μM or 100 μM TCN-P, or 100 μM TCN, followed by heating across 12 temperatures (43–61.7°C). Samples were pooled, clarified, and processed using a modified SP3 cleanup and digestion protocol [45]. Peptides were labeled using TMT16-plex reagents (Thermo Fisher Scientific) and fractionated by HiRIEF prior to LC–MS/MS analysis.

Quantitative proteomics and phosphoproteomics sample preparation was performed as described previously [31, 46]. For phosphoproteomics, phosphopeptides were enriched using IMAC/TiO₂ prior to analysis.

### Quantification of TCN and TCN-P in cell extracts and media

Intracellular and extracellular concentrations of TCN and TCN-P were quantified by LC–MS/MS. For intracellular metabolite analysis, 5 × 10⁶ cells were collected and extracted in 500 μL of ice-cold 90% methanol. Samples were disrupted by bead-based homogenization using a tungsten bead (30 kHz, 2 min), followed by centrifugation at 14,000 × g for 10 min at 4°C. A total of 400 μL of the cleared supernatant was transferred to a 96-well plate and dried under a nitrogen stream. Dried extracts were reconstituted in 100 μL of 0.1% formic acid in water, and 0.5 μL of each sample was injected for LC–MS/MS analysis.

For extracellular measurements, 50 μL of culture medium was mixed with 450 μL of 100% methanol and vortexed for 2 min to precipitate proteins. Samples were centrifuged at 14,000 × g for 10 min at 4°C, and 400 μL of the supernatant was transferred to a 96-well plate and dried under nitrogen. Samples were reconstituted in 100 μL of 0.1% formic acid in water, and 1 μL was injected for LC–MS/MS analysis.

Quantitative analysis was performed using a triple quadrupole mass spectrometer (Agilent 6495D, Muninn platform) operated in multiple reaction monitoring (MRM) mode. Peak integration and quantification were performed using vendor-supplied software. Relative intracellular and extracellular concentrations of TCN and TCN-P were calculated based on extracted ion chromatograms.

### Experimental design and statistical analysis

All proteomics experiments were performed with at least three biological replicates. Statistical analyses were conducted using one-way ANOVA or moderated t-test as appropriate. Differential protein abundance was calculated using DEqMS [47], and differentially expressed proteins were defined using the following thresholds: ALL-SIL, |log₂FC| > 0.5 and adjusted p-value < 0.01; KASUMI-10, |log₂FC| > 0.38 and adjusted p-value < 0.01; P12-ICHIKAWA and KASUMI-2, |log₂FC| > 0.38 and adjusted p-value < 0.05. Phosphoproteomic comparisons were performed using limma [48]. P-values were adjusted for multiple testing using the Benjamini–Hochberg method unless otherwise stated.

### Databases and bioinformatic analyses

Drug sensitivity (DSRT) data were obtained from the FORALL database [32]. PRISM drug repurposing data were obtained from DepMap [33]. Kaplan–Meier survival analysis of AML patients was obtained from the KM plotter [49]. GSEA and ssGSEA analyses were performed using msigdbr and GSVA [50–52]. Pathway enrichment networks were visualized using aPEAR [53], and kinase activity inference was performed using KSEA[54]. Figures were created using BioRender, GraphPad Prism and Adobe Illustrator.

## Data availability

The mass spectrometry proteomics data will be deposited to the ProteomeXchange Consortium via the PRIDE partner repository.

## Acknowledgements

This study was supported by grants from the Swedish Childhood Cancer Foundation (R.J., TJ2016-0035, PR2016-0019, PR2019-0025, PR2022-0009 and PR2024-0065); The Swedish Cancer Society (R.J., 23 2843 Pj and 24 0866 SIA); the Swedish Research Council (R.J., 2017-01653); the Felix Mindus Contribution to Leukemia Research (R.J.); the Dr. Åke Olsson Foundation for Hematological Research (R.J., 2021-00130); Cancer Research Foundations of Radiumhemmet (R.J., 221132 and 241243); and KI Research Foundation Grants (X.Q., 2024-02353); R.J. acknowledges the Karolinska Institutet and Science for Life Laboratory. The authors would like to thank Owen Clemente Mc Swiney and Högni Fjalarsson for their assistance during the project. Figures 2B, 4A, 4D, 6D, and Supplementary Figures 3A and 5E were created in BioRender. Proteomics, L. (2026) https://BioRender.com/52n4zye.

## References

1. Malard, F. and M. Mohty, Acute lymphoblastic leukaemia. Lancet, 2020. 395(10230): p. 1146–1162.

2. Pagliaro, L., et al., Acute lymphoblastic leukaemia. Nat Rev Dis Primers, 2024. 10(1): p. 41.

3. Wang, J.Y., et al., Metabolomic insights into pathogenesis and therapeutic potential in adult acute lymphoblastic leukemia. Proc Natl Acad Sci U S A, 2025. 122(7): p. e2423169122.

4. Mullen, N.J. and P.K. Singh, Nucleotide metabolism: a pan-cancer metabolic dependency. Nat Rev Cancer, 2023. 23(5): p. 275–294.

5. Chandel, N.S., Nucleotide Metabolism. Cold Spring Harb Perspect Biol, 2021. 13(7).

6. Tran, D.H., et al., De novo and salvage purine synthesis pathways across tissues and tumors. Cell, 2024. 187(14): p. 3602–3618 e20.

7. Toksvang, L.N., et al., Maintenance therapy for acute lymphoblastic leukemia: basic science and clinical translations. Leukemia, 2022. 36(7): p. 1749–1758.

8. Galmarini, C.M., J.R. Mackey, and C. Dumontet, Nucleoside analogues and nucleobases in cancer treatment. Lancet Oncol, 2002. 3(7): p. 415–24.

9. Rudd, S.G., Targeting pan-essential pathways in cancer with cytotoxic chemotherapy: challenges and opportunities. Cancer Chemother Pharmacol, 2023. 92(4): p. 241–251.

10. Wotring, L.L., et al., Mechanism of activation of triciribine phosphate (TCN-P) as a prodrug form of TCN. Cancer Treat Rep, 1986. 70(4): p. 491–7.

11. Plagemann, P.G., Transport, phosphorylation, and toxicity of a tricyclic nucleoside in cultured Novikoff rat hepatoma cells and other cell lines and relase of its monophosphate by the cells. J Natl Cancer Inst, 1976. 57(6): p. 1283–95.

12. Moore, E.C., R.B. Hurlbert, and S.P. Massia, Inhibition of CCRF-CEM human leukemic lymphoblasts by triciribine (tricyclic nucleoside, TCN, NSC-154020). Accumulation of drug in cells and comparison of effects on viability, protein synthesis and purine synthesis. Biochem Pharmacol, 1989. 38(22): p. 4037–44.

13. Ptak, R.G., et al., Phosphorylation of triciribine is necessary for activity against HIV type 1. AIDS Res Hum Retroviruses, 1998. 14(15): p. 1315–22.

14. Frejno, M., et al., Proteome activity landscapes of tumor cell lines determine drug responses. Nat Commun, 2020. 11(1): p. 3639.

15. Shedden, K., et al., A rational approach to personalized anticancer therapy: chemoinformatic analysis reveals mechanistic gene-drug associations. Pharm Res, 2003. 20(6): p. 843–7.

16. Yang, L., et al., Akt/protein kinase B signaling inhibitor-2, a selective small molecule inhibitor of Akt signaling with antitumor activity in cancer cells overexpressing Akt. Cancer Res, 2004. 64(13): p. 4394–9.

17. Berndt, N., et al., The Akt activation inhibitor TCN-P inhibits Akt phosphorylation by binding to the PH domain of Akt and blocking its recruitment to the plasma membrane. Cell Death Di]er, 2010. 17(11): p. 1795–804.

18. Shelton, J., et al., Metabolism, Biochemical Actions, and Chemical Synthesis of Anticancer Nucleosides, Nucleotides, and Base Analogs. Chem Rev, 2016. 116(23): p. 14379–14455.

19. Lu, C.H., et al., Preclinical testing of clinically applicable strategies for overcoming trastuzumab resistance caused by PTEN deficiency. Clin Cancer Res, 2007. 13(19): p. 5883–8.

20. Wotring, L.L., et al., Dual mechanisms of inhibition of DNA synthesis by triciribine. Cancer Res, 1990. 50(16): p. 4891–9.

21. Moore, E.C., et al., Inhibition of two enzymes in de novo purine nucleotide synthesis by triciribine phosphate (TCN-P). Biochem Pharmacol, 1989. 38(22): p. 4045–51.

22. Takahashi, S., Signaling effect, combinations, and clinical applications of triciribine. J Chemother, 2025. 37(6): p. 494–502.

23. Sampath, D., et al., Phase I clinical, pharmacokinetic, and pharmacodynamic study of the Akt-inhibitor triciribine phosphate monohydrate in patients with advanced hematologic malignancies. Leuk Res, 2013. 37(11): p. 1461–7.

24. Aebersold, R. and M. Mann, Mass-spectrometric exploration of proteome structure and function. Nature, 2016. 537(7620): p. 347–55.

25. Leo, I.R., et al., Functional proteoform group deconvolution reveals a broader spectrum of ibrutinib off-targets. Nat Commun, 2025. 16(1): p. 1948.

26. Savitski, M.M., et al., Tracking cancer drugs in living cells by thermal profiling of the proteome. Science, 2014. 346(6205): p. 1255784.

27. Kurzawa, N., et al., Deep thermal profiling for detection of functional proteoform groups. Nat Chem Biol, 2023. 19(8): p. 962–971.

28. Martinez Molina, D., et al., Monitoring drug target engagement in cells and tissues using the cellular thermal shift assay. Science, 2013. 341(6141): p. 84–7.

29. Gaetani, M., et al., Proteome Integral Solubility Alteration: A High-Throughput Proteomics Assay for Target Deconvolution. J Proteome Res, 2019. 18(11): p. 4027–4037.

30. Branca, R.M., et al., HiRIEF LC-MS enables deep proteome coverage and unbiased proteogenomics. Nat Methods, 2014. 11(1): p. 59–62.

31. Leo, I.R., et al., Integrative multi-omics and drug response profiling of childhood acute lymphoblastic leukemia cell lines. Nat Commun, 2022. 13(1): p. 1691.

32. Aswad, L. and R. Jafari, FORALL: an interactive shiny/R web portal to navigate multiomics high-throughput data of pediatric acute lymphoblastic leukemia. Bioinform Adv, 2023. 3(1): p. vbad143.

33. Corsello, S.M., et al., Discovering the anti-cancer potential of non-oncology drugs by systematic viability profiling. Nat Cancer, 2020. 1(2): p. 235–248.

34. Palomero, T., et al., Mutational loss of PTEN induces resistance to NOTCH1 inhibition in T-cell leukemia. Nat Med, 2007. 13(10): p. 1203–10.

35. Hernandez-Armenta, C., et al., Benchmarking substrate-based kinase activity inference using phosphoproteomic data. Bioinformatics, 2017. 33(12): p. 1845–1851.

36. Lei, M., The MCM complex: its role in DNA replication and implications for cancer therapy. Curr Cancer Drug Targets, 2005. 5(5): p. 365–80.

37. Ceccaldi, R. and P. Cejka, Mechanisms and regulation of DNA end resection in the maintenance of genome stability. Nat Rev Mol Cell Biol, 2025. 26(8): p. 586–599.

38. Plagemann, P.G. and J. Erbe, Exit transport of a cyclic nucleotide from mouse L-cells. J Biol Chem, 1977. 252(6): p. 2010–6.

39. Yan, X. and C. Liu, The ATF4-glutamine axis: a central node in cancer metabolism, stress adaptation, and therapeutic targeting. Cell Death Discov, 2025. 11(1): p. 390.

40. Ben-Sahra, I., et al., Sestrin2 integrates Akt and mTOR signaling to protect cells against energetic stress-induced death. Cell Death Di]er, 2013. 20(4): p. 611–9.

41. Struyf, N., et al., Delineating functional and molecular impact of ex vivo sample handling in precision medicine. NPJ Precis Oncol, 2024. 8(1): p. 38.

42. Erkers, T., et al., Data-Driven Molecular Hallmarks of Acute Myeloid Leukemia: Biological, Prognostic and Therapeutic Implications. Blood, 2022. 140(Supplement 1): p. 6299–6299.

43. Osa-Andrews, B., et al., Development of Novel Intramolecular FRET-Based ABC Transporter Biosensors to Identify New Substrates and Modulators. Pharmaceutics, 2018. 10(4).

44. Goroshchuk, O., et al., Thermal proteome profiling identifies PIP4K2A and ZADH2 as off-targets of Polo-like kinase 1 inhibitor volasertib. FASEB J, 2021. 35(7): p. e21741.

45. Moggridge, S., et al., Extending the Compatibility of the SP3 Paramagnetic Bead Processing Approach for Proteomics. J Proteome Res, 2018. 17(4): p. 1730–1740.

46. Tape, C.J., et al., Reproducible automated phosphopeptide enrichment using magnetic TiO2 and Ti-IMAC. Anal Chem, 2014. 86(20): p. 10296–302.

47. Zhu, Y., et al., DEqMS: A Method for Accurate Variance Estimation in Differential Protein Expression Analysis. Mol Cell Proteomics, 2020. 19(6): p. 1047–1057.

48. Smyth, G.K., Linear models and empirical bayes methods for assessing differential expression in microarray experiments. Stat Appl Genet Mol Biol, 2004. 3: p. Article3.

49. Gyor]y, B., Discovery and ranking of the most robust prognostic biomarkers in serous ovarian cancer. Geroscience, 2023. 45(3): p. 1889–1898.

50. Subramanian, A., et al., Gene set enrichment analysis: a knowledge-based approach for interpreting genome-wide expression profiles. Proc Natl Acad Sci U S A, 2005. 102(43): p. 15545–50.

51. Liberzon, A., et al., Molecular signatures database (MSigDB) 3.0. Bioinformatics, 2011. 27(12): p. 1739–40.

52. Hanzelmann, S., R. Castelo, and J. Guinney, GSVA: gene set variation analysis for microarray and RNA-seq data. BMC Bioinformatics, 2013. 14: p. 7.

53. Kerseviciute, I. and J. Gordevicius, aPEAR: an R package for autonomous visualization of pathway enrichment networks. Bioinformatics, 2023. 39(11).

54. Wiredja, D.D., M. Koyuturk, and M.R. Chance, The KSEA App: a web-based tool for kinase activity inference from quantitative phosphoproteomics. Bioinformatics, 2017. 33(21): p. 3489–3491.

