## Supplementary figures for "Proteomic profiling reveals pleiotropic antimetabolite activity of triciribine in acute lymphoblastic leukemia"

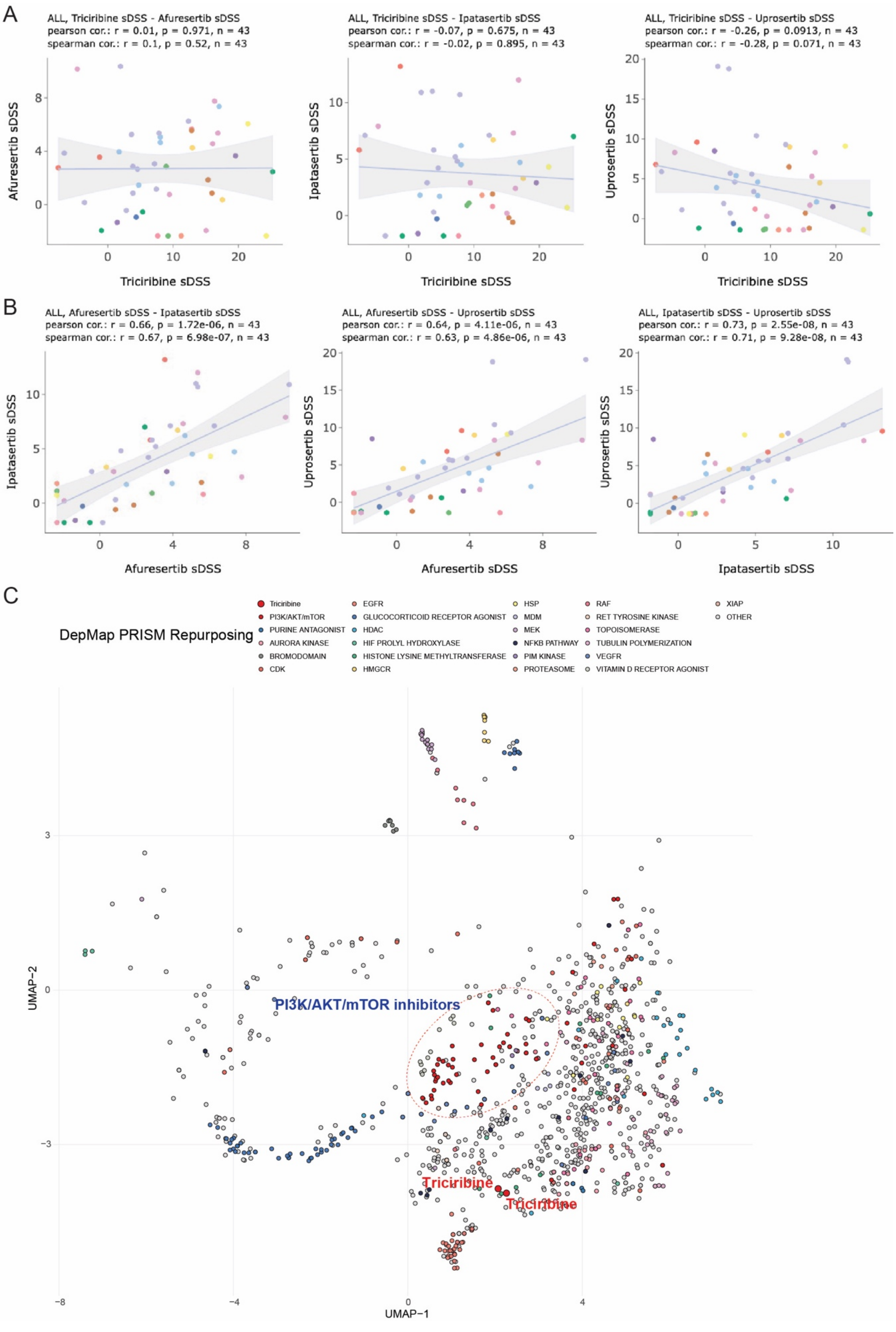
A
ALL, Triciribine sDSS - Afuresertib sDSS
pearson cor.: r = 0.01, p = 0.971, n = 43
spearman cor.: r = 0.1, p = 0.52, n = 43
Afuresertib sDSS
Triciribine sDSS
ALL, Triciribine sDSS - Ipatasertib sDSS
pearson cor.: r = -0.07, p = 0.675, n = 43
spearman cor.: r = -0.02, p = 0.895, n = 43
Ipatasertib sDSS
Triciribine sDSS
ALL, Triciribine sDSS - Uprosertib sDSS
pearson cor.: r = -0.26, p = 0.0913, n = 43
spearman cor.: r = -0.28, p = 0.071, n = 43
Uprosertib sDSS
Triciribine sDSS
B
ALL, Afuresertib sDSS - Ipatasertib sDSS
pearson cor.: r = 0.66, p = 1.72e-06, n = 43
spearman cor.: r = 0.67, p = 6.98e-07, n = 43
Ipatasertib sDSS
Afuresertib sDSS
ALL, Afuresertib sDSS - Uprosertib sDSS
pearson cor.: r = 0.64, p = 4.11e-06, n = 43
spearman cor.: r = 0.63, p = 4.86e-06, n = 43
Uprosertib sDSS
Afuresertib sDSS
ALL, Ipatasertib sDSS - Uprosertib sDSS
pearson cor.: r = 0.73, p = 2.55e-08, n = 43
spearman cor.: r = 0.71, p = 9.28e-08, n = 43
Uprosertib sDSS
Ipatasertib sDSS
C
DepMap PRISM Repurposing
Triciribine
PI3K/AKT/mTOR
PURINE ANTAGONIST
AURORA KINASE
BROMODOMAIN
CDK
EGFR
GLUCOCORTICOID RECEPTOR AGONIST
HDAC
HIF PROLYL HYDROXYLASE
HISTONE LYSINE METHYLTRANSFERASE
HMGCR
HSP
MDM
MEK
NFKB PATHWAY
PIM KINASE
PROTEASOME
RAF
RET TYROSINE KINASE
TOPOISOMERASE
TUBULIN POLYMERIZATION
VEGFR
VITAMIN D RECEPTOR AGONIST
XIAP
OTHER
PI3K/AKT/mTOR inhibitors
Triciribine
Triciribine
UMAP−2
UMAP−1

**Supplementary Figure 1. Triciribine exhibits a distinct drug sensitivity profile compared with Akt inhibitors.**

**A**, Pairwise correlation analysis comparing the drug sensitivity profile of triciribine with afuresertib, ipatasertib, and uprosertib across 43 ALL cell lines.

**B**, Pairwise correlation analysis of drug sensitivity profiles among afuresertib, ipatasertib, and uprosertib across the same ALL cell line panel.

**C**, Two-dimensional UMAP projection of drug sensitivity profiles from the PRISM repurposing dataset. Compounds are positioned based on cosine similarity of killing profiles, and compounds with shared annotated mechanisms of action are indicated by color.

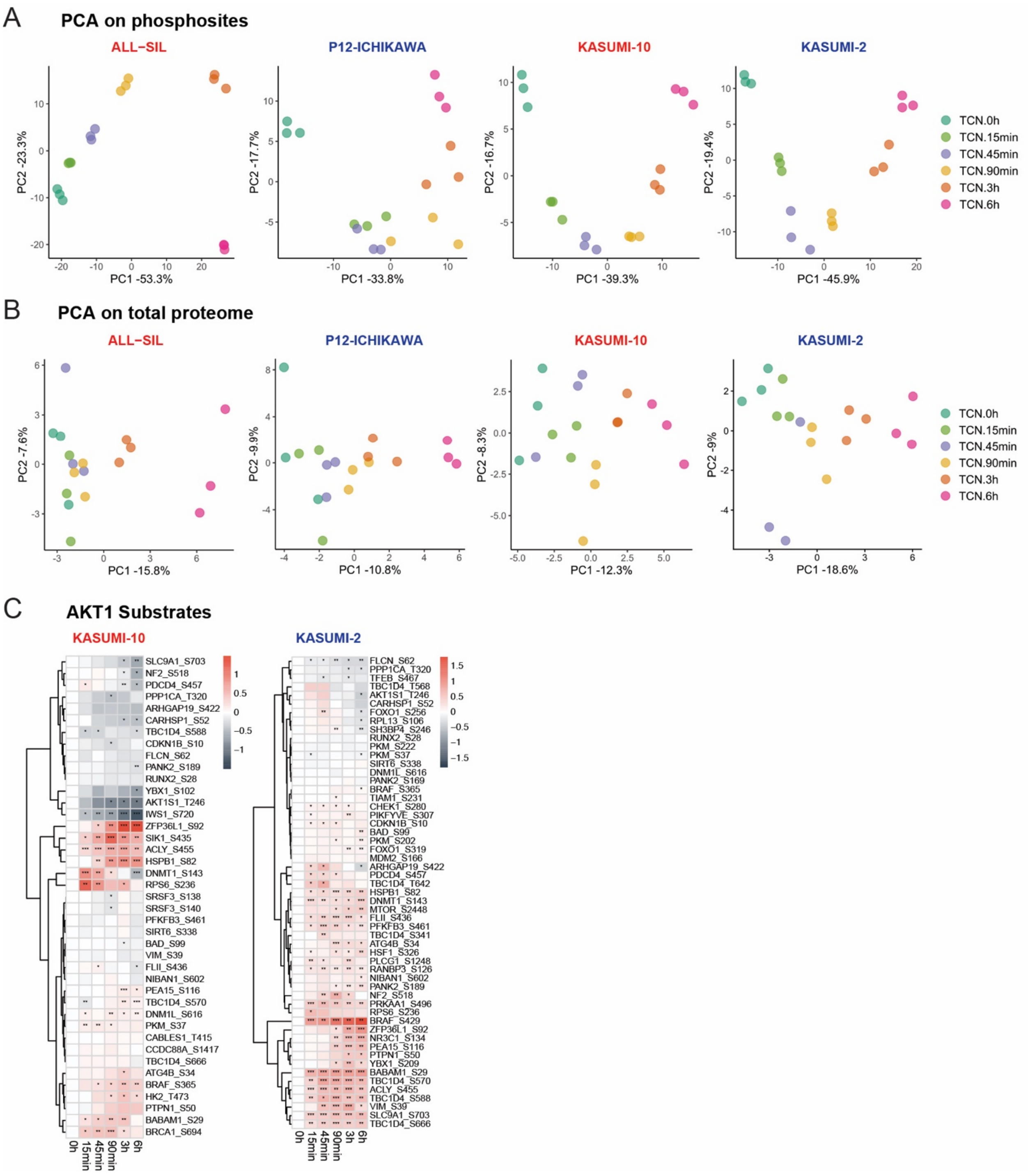


**Supplementary Figure 2. Global phosphoproteomic and proteomic profiling following triciribine treatment.**

**A**, Principal component analysis (PCA) of phosphosite-level quantitative data from four ALL cell lines treated with triciribine across indicated time points.

**B**, PCA of total proteome quantitative data from the same samples.

**C**, Phosphorylation changes of known Akt1 substrates across time points relative to 0 h in KASUMI-10 and KASUMI-2 cells.

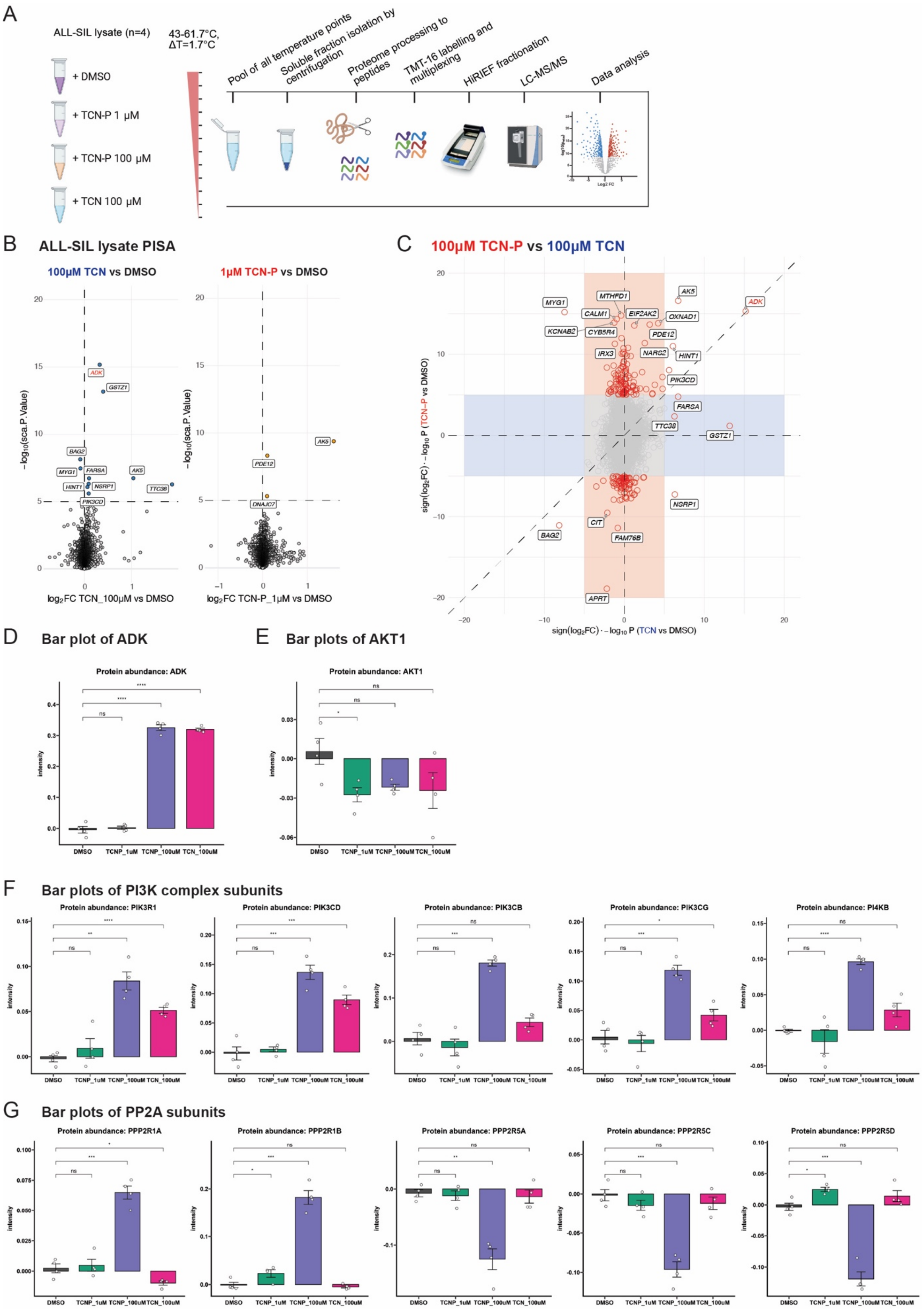

A
ALL-SIL lysate (n=4)
+ DMSO
+ TCN-P 1 µM
+ TCN-P 100 µM
+ TCN 100 µM
43-61.7°C, ΔT=1.7°C
Pool of all temperature points
Soluble fraction isolation by centrifugation
Proteome processing to peptides
TMT-16 labelling and multiplexing
HiRIEF fractionation
LC-MS/MS
Data analysis
B
ALL-SIL lysate PISA
100µM TCN vs DMSO
1µM TCN-P vs DMSO
−log10(sca.P.Value)
log2FC TCN_100µM vs DMSO
log2FC TCN-P_1µM vs DMSO
ADK
GSTZ1
BAG2
MYG1
FARSA
AK5
HINT1
NSRP1
TTC38
PIK3CD
PDE12
DNAJC7
C
100µM TCN-P vs 100µM TCN
sign(log2FC) · −log10 P (TCN-P vs DMSO)
sign(log2FC) · −log10 P (TCN vs DMSO)
MYG1
MTHFD1
AK5
ADK
CALM1
EIF2AK2
OXNAD1
KCNAB2
CYB5R4
PDE12
IRX3
NARS2
HINT1
PIK3CD
FARSA
TTC38
GSTZ1
NSRP1
CIT
BAG2
FAM76B
APRT
D
Bar plot of ADK
Protein abundance: ADK
E
Bar plots of AKT1
Protein abundance: AKT1
F
Bar plots of PI3K complex subunits
Protein abundance: PIK3R1
Protein abundance: PIK3CD
Protein abundance: PIK3CB
Protein abundance: PIK3CG
Protein abundance: PI4KB
G
Bar plots of PP2A subunits
Protein abundance: PPP2R1A
Protein abundance: PPP2R1B
Protein abundance: PPP2R5A
Protein abundance: PPP2R5C
Protein abundance: PPP2R5D
intensity
DMSO
TCNP_1uM
TCNP_100uM
TCN_100uM

**Supplementary Figure 3. Expanded PISA analysis of candidate triciribine-interacting proteins.**

**A**, Experimental design and workflow of the Proteome Integral Solubility Alteration (PISA) assay.

**B**, Volcano plots of PISA results showing changes in protein solubility following treatment with 100 μM triciribine or 1 μM TCN-P compared with DMSO control.

**C**, Comparison of PISA results between 100 μM TCN-P and 100 μM triciribine treatments. The x- and y-axes represent signed −log10(p-values) multiplied by the direction of solubility change.

**D–G**, Bar plots showing solubility changes of selected candidate TCN-P–interacting proteins identified by PISA.

A **ALL−SIL TCN 24h: GO BP up-regulation**

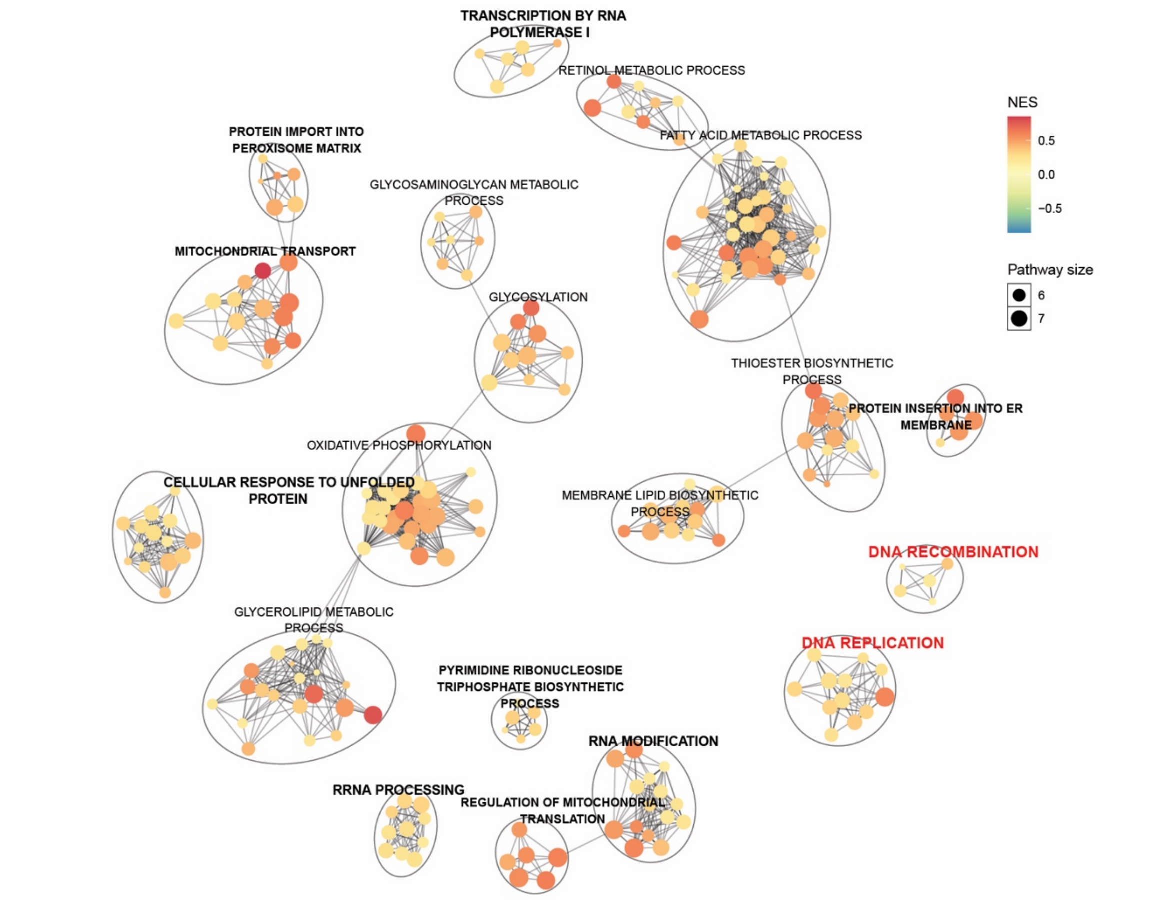


B **ALL−SIL TCN 24h: GSEA**

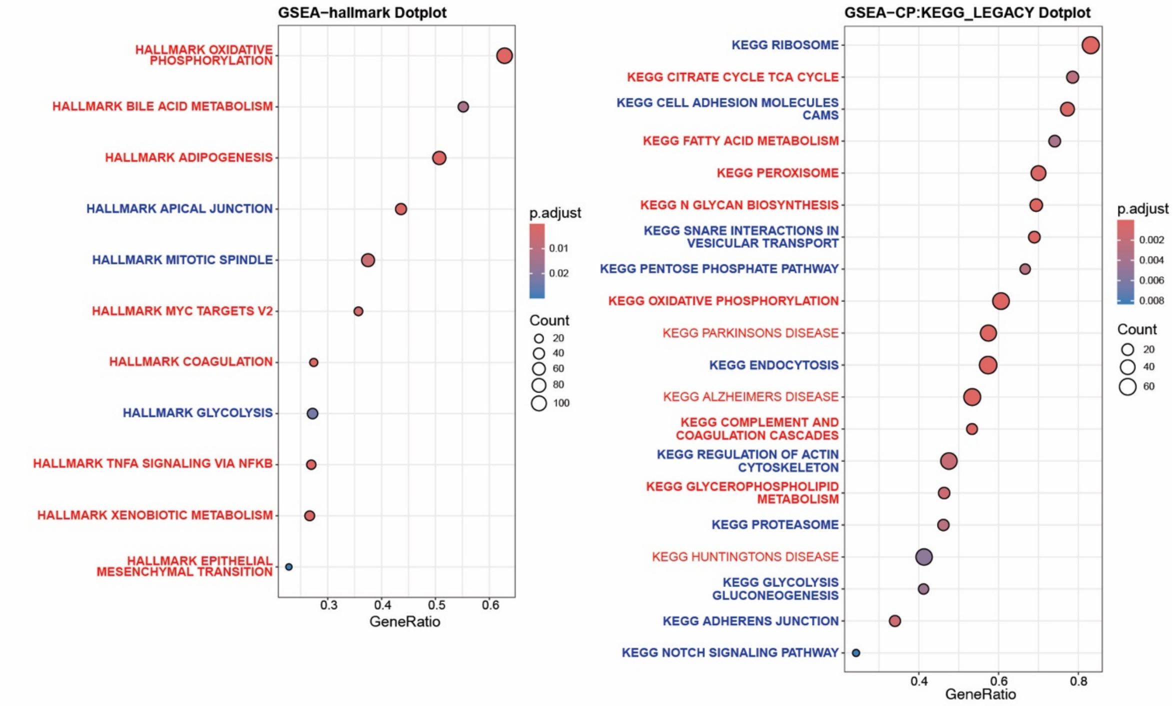

**Supplementary Figure 4. Pathway-level proteomic responses.**

**A**, aPEAR visualization of significantly up-regulated biological process networks identified by ssGSEA in ALL-SIL cells treated with triciribine for 24 h compared with DMSO. **B**, Dot plot summarizing gene set enrichment analysis results using Hallmark and KEGG databases for ALL-SIL cells treated with triciribine for 24 h compared with DMSO. Red text indicates pathways or signatures upregulated in triciribine-treated cells relative to DMSO, whereas blue text indicates downregulated pathways.

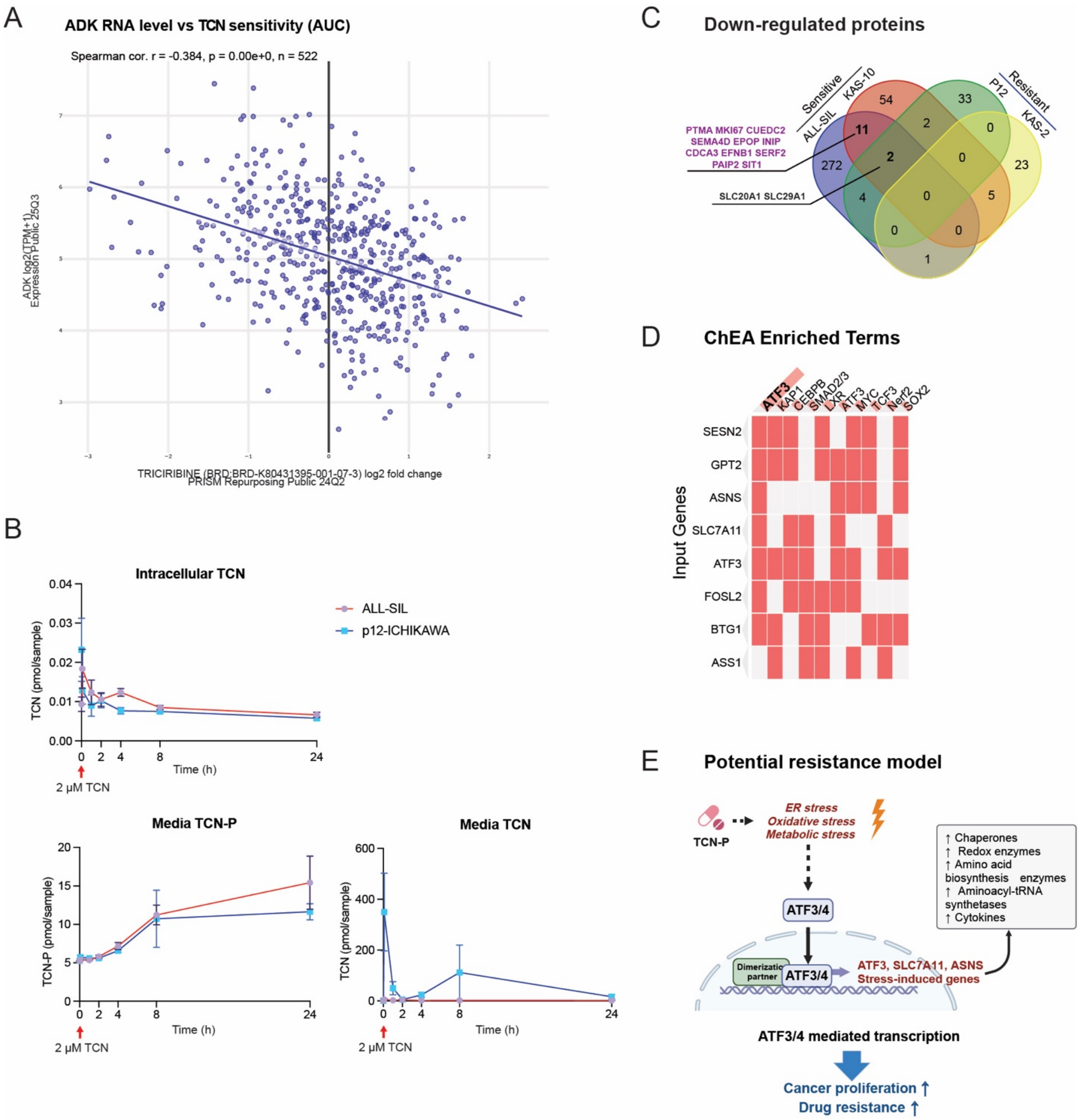


**Supplementary Figure 5. ADK expression, triciribine metabolism, and comparative proteomic responses.**

**A**, Correlation between ADK mRNA expression and triciribine sensitivity in the PRISM repurposing cohort.

**B**, Time-course quantification of intracellular triciribine, extracellular triciribine monophosphate (TCN-P), and extracellular triciribine concentrations following treatment with 2 μM triciribine in ALL-SIL and P12-ICHIKAWA cells.

**C**, Venn diagram showing the overlap of proteins significantly downregulated by triciribine treatment at both 6 h and 24 h across four ALL cell lines.
**D**, ChEA analysis identifying candidate transcription factors associated with proteins upregulated in less sensitive ALL cell lines following triciribine treatment.
**E**, Schematic representation of a proposed model summarizing cellular features associated with reduced triciribine sensitivity in ALL cells.

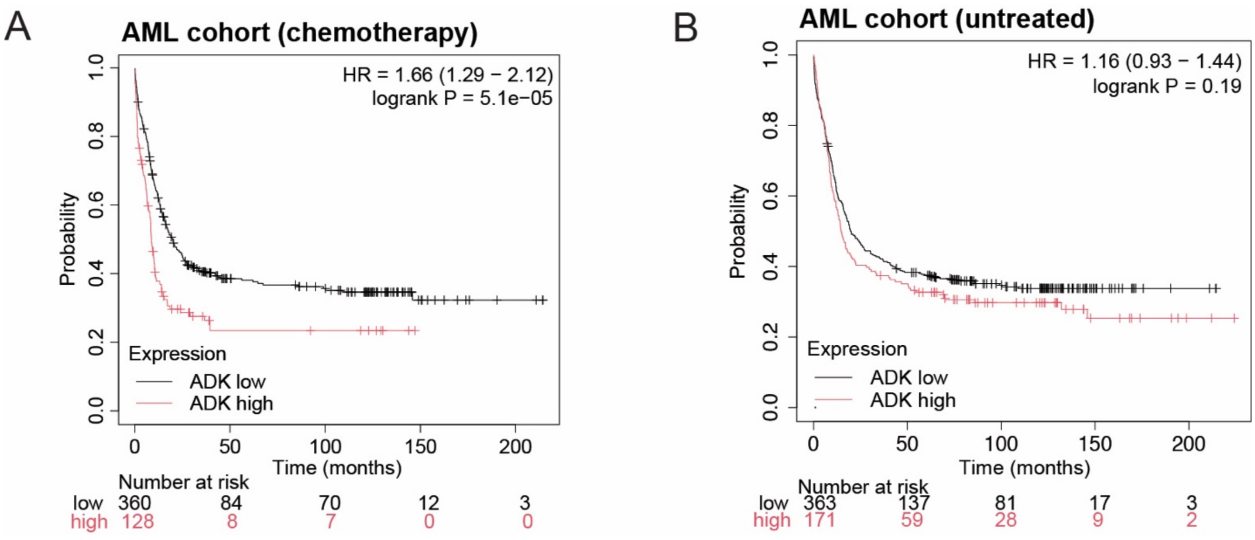


**Supplementary Figure 6. A-B**, Kaplan–Meier survival analysis of chemotherapy-treated or untreated AML patients stratified by ADK expression.
